# Identification of large-scale genomic rearrangements during wheat evolution and the underlying mechanisms

**DOI:** 10.1101/478933

**Authors:** Inbar Bariah, Danielle Keidar-Friedman, Khalil Kashkush

## Abstract

Following allopolyploidization, nascent polyploid wheat species react with massive genomic rearrangements, including deletion of transposable element-containing sequences. While such massive rearrangements are considered to be a prominent process in wheat genome evolution and speciation, their structure, extent, and underlying mechanisms remain poorly understood. In this study, we retrieved ~3500 insertions of a specific variant of *Fatima*, one of the most dynamic long-terminal repeat retrotransposons in wheat from the recently available high-quality genome drafts of *Triticum aestivum* (bread wheat) and *Triticum turgidum ssp. dicoccoides* or wild emmer, the allotetraploid mother of all modern wheats. The dynamic nature of *Fatima* facilitated the identification of large (i.e., up to ~ 1 million bases) *Fatima*-containing insertions/deletions (InDels) upon comparison of bread wheat and wild emmer genomes. We characterized 11 such InDels using computer-assisted analysis followed by PCR validation, and found that they occurred via unequal intra-strand recombination or double-strand break events. In most cases, InDels breakpoints were located within transposable element sequences. Additionally, we observed one case of introgression of novel DNA fragments from an unknown source into the wheat genome. Our data thus indicate that massive large-scale DNA rearrangements might play a prominent role in wheat speciation.

## Introduction

The evolution of pasta and bread wheats (the *Triticum-Aegilops* group) involved two separate allopolyploidization events. The first occurred ~0.5 MYA and included the hybridization of *Triticum urartu* (donor of the A genome) and an unknown species from section *Sitopsis* (donor of the B genome), leading to the formation of the allotetraploid wild emmer *T. turgidum* ssp*. dicoccoides* (genome AABB) (Dvořák et al. 1993; Feldman and Levy 2005; Ling et al. 2018). The initial domestication of the wild emmer gave rise to the domesticated emmer wheat *T. turgidum* ssp*. dicoccum* (genome AABB), followed by selection of free-threshing durum wheat (*T. turgidum* ssp*. durum*, genome AABB) (Avni et al. 2017). The second allopolyploidization event that occurred ~10,000 years ago included hybridization of the domesticated emmer and *Aegilops tauschii* (donor of the D genome) and led to the generation of the bread wheat *T. aestivum* (genome AABBDD) (Feldman and Levy 2005; Petersen et al. 2006).

Wheat allopolyploids are relatively young species and thus are expected to show limited genetic variation due to the “polyploidy diversity bottleneck”. This diversity bottleneck is the result of several factors, namely the short time since allopolyploid formation which is insufficient for the accumulation of mutations, the involvement of only few individuals from the progenitor species in the allopolyploidization event and reproductive isolation of the newly formed allopolyploid from the parental species (Stebbins 1950; Feldman and Levy 2012). Nevertheless, wheat allopolyploids show wider morphological variation, occupy a greater diversity of ecological niches and proliferate over larger geographical areas, relative to their diploid ancestors (Feldman and Levy 2012). Indeed, the accelerated genome evolution triggered by allopolyploidy may be largely responsible for the wide genetic and morphologic diversity observed in wheat allopolyploids.

Allopolyploidy was shown to trigger a series of revolutionary (i.e., occurring immediately after allopolyploidization) as well evolutionary (i.e., occurring during the life of the allopolyploid species) genomic changes in wheat allopolyploids, which might not be attainable at the diploid level (Feldman and Levy 2005; Feldman and Levy 2012). These genomic changes can include the activation of transposable elements (TEs), together with massive and reproducible elimination of TE-containing sequences, as was reported for newly formed wheat allopolyploids (Shaked et al. 2001; Kraitshtein et al. 2010; Yaakov and Kashkush 2012; Ben-David et al. 2013; Yaakov et al. 2013). TEs, corresponding to fragments of DNA able to “move” and proliferate within the host genome, account for over 80% of the wheat genome (Charles et al. 2008; Consortium 2014; Avni et al. 2017; Clavijo et al. 2017; Appels et al. 2018). The majority of TEs in wheat allopolyploid genomes are derived from long-terminal repeat retrotransposons (LTRs) that contribute to the highly repetitive nature of those genomes (Avni et al. 2017; Clavijo et al. 2017; Appels et al. 2018). Due to their highly repetitive nature, TEs can interact in a disruptive manner during both meiotic recombination and DNA repair processes, leading to a variety of genomic rearrangements, including sequence translocations, duplications and elimination (Devos et al. 2002; Ma et al. 2004; Hedges and Deininger 2007; Kraitshtein et al. 2010; Fedoroff 2012).

The mechanism(s) of DNA sequence elimination, including deletion of TE-containing sequences following allopolyploidization events, has yet to be identified. In this study, a specific variant of *Fatima*, a well-represented family of *gypsy* LTR retrotransposons, was used to identify flanking DNA sequences that had been eliminated from wheat allopolyploid genomes. In addition, InDel (insertion/deletion) breakpoints were identified and further characterized. Detailed analysis of 11 InDels gave rise to possible mechanisms involved in DNA rearrangements following allopolyploidization and/or domestication processes. Finally, the possible role of DNA rearrangements in speciation and domestication is discussed.

## Results and Discussion

### Addressing *Fatima* LTR retrotransposons to identify large-scale sequence variations between wild emmer and bread wheat

In a previous study, we reported a significant decrease in relative copy numbers of *Fatima* elements in newly formed allohexaploids, relative to the expected additive parental copy number (Yaakov et al. 2013). A possible explanation for this result was the rapid elimination of *Fatima*-containing sequences following allopolyploidization events. This, together with the availability of genome drafts for various wheat species, facilitated the identification of large-scale genomic rearrangements between wild emmer and bread wheat.

The consensus sequence of the autonomous *Fatima* element was used as a query in a search using MAK software designed to retrieve *Fatima* insertions, together with their flanking sequences (500 bp from each side), from the draft genomes of wild emmer and bread wheat. Overall, 1,761 intact *Fatima* insertions were retrieved from the wild emmer genome and 1,741 intact *Fatima* insertions were retrieved from the bread wheat genome. The majority of retrieved *Fatima* insertions (97.4% in wild emmer and 97.6% in bread wheat) were located within the B sub-genome (Supplemental Fig. S1). The remaining retrieved *Fatima* insertions were found in the A sub-genome (36 insertions in wild emmer and 33 insertions in bread wheat), or were unmapped (10 insertions in wild emmer and 8 insertions in bread wheat). The B sub-genome specificity of specific *Fatima* variants in polyploid and diploid wheat species was also reported in previous studies (Salina et al. 2011; Yaakov et al. 2013; Wicker et al.2018).

The wheat B sub-genome may have undergone massive modifications (yielding the differential genome), as the BB genome donor has yet to be identified and A and D sub-genomes are conserved (termed the pivotal genome), a phenomenon referred to as ‘pivotal-differential’ genome evolution (Mirzaghaderi and Mason 2017). Thus, the B sub-genome was a promising target in efforts aimed at identifying large-scale genomic rearrangements. In this study, we accordingly focused specifically on chromosomes 3B and 5B, given how the first notable high-quality sequence assembly of wheat was reported for bread wheat chromosome 3B (Paux et al. 2008), the largest chromosome in the wild emmer and bread wheat genomes. In addition, chromosome 5B contains the major chromosome pairing *Ph1* locus (Feldman and Levy 2012). In wild emmer, 268 *Fatima* insertions were retrieved from chromosome 3B and 274 *Fatima* insertions were retrieved from chromosome 5B, while in bread wheat, 274 *Fatima* insertions were retrieved from chromosome 3B and 277 *Fatima* insertions were retrieved from chromosome 5B. Comparative analysis revealed that while the majority of *Fatima* insertions in chromosomes 3B and 5B are common to wild emmer and bread wheat (i.e., monomorphic insertions), ~15% of the insertions occurred at polymorphic insertion sites. Several sources for such polymorphism were identified. In ~5% of the cases, the presence (i.e., full sites) vs. the absence (i.e., empty sites) of *Fatima* with notable target site duplications (TSDs) were noted. In ~57% of the cases, insertions and/or deletions were detected within the *Fatima* element; in some of these instances, the deletion also included part of the *Fatima*-flanking (i.e., chimeric) sequences. In 34% of the cases, large-scale rearrangements of *Fatima*-containing sequences ranging in size from 13 kb to 4.4 Mb), including large scale deletions, introgressions, and duplications, were seen. Finally, because of assembly artefacts, some 4% of the readings were false positives. Deletions and other rearrangements are known to be prevalent among LTR retrotransposon elements and retrotransposon-containing sequences (Devos et al. 2002; Ma et al. 2004; Bennetzen et al. 2005; Kraitshtein et al. 2010). Here, addressing *Fatima*, a well-represented *gypsy* LTR retrotransposon family in wheat, facilitated the identification of such large-scale genomic rearrangements between wild emmer and bread wheat.

Detailed analysis of 11 cases of large-scale rearrangements using a chromosome walking approach and dot plot sequence alignments (Supplemental Fig. S2) of the affected loci in the wild emmer and bread wheat genomes revealed 9 instances of long deletions in bread wheat (5 in chromosome 3B and 4 in chromosome 5B), the introduction of a new DNA fragment, and a single example of copy number variation of a long tandem repeat in chromosome 5B. In all 11 cases, InDel breakpoints were identified as the borders between high sequence similarity regions (i.e., 95% sequence identity or higher for a word size of 100) to regions that showed no sequence similarity (i.e., lower than 95% sequence identity for a word size of 100) using dot plot representations of the sequence alignments between the orthologous loci in the wild emmer and bread wheat genomes. The lengths of the eliminated and/or introduced sequences were defined as the distances between the 5’ and the 3’ breakpoints. Table 1 summarizes the *in silico* characterization of the 11 loci in wild emmer vs. bread wheat. Note that although *Fatima*-containing sequences were found to be eliminated from the bread wheat genome, the total number of retrieved *Fatima* insertions was similar in wild emmer and bread wheat, suggesting that *Fatima* was most likely activated following allohexaploidization, leading to the existence of new *Fatima* insertions in the bread wheat genome. A similar pattern was described for a terminal-repeat retrotransposon in miniature (TRIM) family termed *Veju* in the first four generations of a newly formed wheat allohexaploid (Kraitshtein et al. 2010).

**Table 1.** *In silico* characterization of large-sequence variations identified in chromosomes 3B and 5B in the bread wheat vs. wild emmer genomes.

| Locus<sup>1</sup><br>ID | Location |  | Locus length (bp)<sup>4</sup> |  | Type of rearrangement |
| --- | --- | --- | --- | --- | --- |
|  | Wild emmer<sup>2</sup> | Bread wheat<sup>3</sup> | Wild emmer | Bread wheat |  |
| **5B<sub>1</sub>** | 5B:566939353-567135082 | 5B:561057394-561064945 | 195730 | 7552 | deletion in bread wheat |
| **3B<sub>1</sub>** | 3B:774200469-774452950 | 3B:760803787-760805537 | 252482 | 1751 | deletion in bread wheat |
| **5B<sub>2</sub>** | 5B:516702290-516721374 | 5B:511383608-511385023 | 19085 | 1416 | deletion in bread wheat |
| **5B<sub>3</sub>** | 5B:363487255-363551431 | 5B:349934346-349934349 | 64177 | 4 | deletion in bread wheat |
| **5B<sub>4</sub>** | 5B:587350647-587364130 | 5B:581381548-581381551 | 13484 | 4 | deletion in bread wheat |
| **3B<sub>2</sub>** | 3B:284755035-284771490 | 3B:286353814-286353819 | 16456 | 6 | deletion in bread wheat |
| **3B<sub>3</sub>** | 3B:493386824-493410158 | 3B:482234389-482234390 | 23335 | 2 | deletion in bread wheat |
| **3B<sub>4</sub>** | 3B:538946011-540047920 | 3B:527682008-527682029 | 1101910 | 22 | deletion in bread wheat |
| **3B<sub>5</sub>** | 3B:606914695-606946995 | 3B:596314588-596314620 | 32301 | 33 | deletion in bread wheat |
| **5B<sub>5</sub>** | 5B:610009239-610050693 | 5B:603942312-603952982 | 41455 | 10671 | introgression of new DNA fragment |
| **5B<sub>6</sub>** | 5B:84661892-85585936 | 5B:81624361-82045980 | 924045 | 421620 | copy number variation |

<sup>1</sup> The first number and letter in the locus ID refer to the chromosome in which the genomic locus is found
<sup>2</sup> WEWSeq\_v.1.0 (http://wewseq.wix.com/consortium) coordinates.
<sup>3</sup> IWGSC ([downloaded in June 2017 from: http://plants.ensembl.org/Triticum\_aestivum/Info/Index](http://plants.ensembl.org/Triticum_aestivum/Info/Index)) coordinates.
<sup>4</sup> Locus length was determined as the genetic distance between the 5' and 3' breakpoints/borders of the sequence variation identified using dot plot alignment between the wild emmer and bread wheat genomes (minimum repeat length of 100 bp and 95% repeat identity; Supplemental Fig. S2). For locus 5B<sub>6</sub>, the borders of the repeat units in wild emmer were identified based on dot plot comparison of the locus surrounding locus 5B<sub>6</sub> in wild emmer against its self (minimum repeat length of 100 bp and 95% repeat identity; Supplemental Fig. S3C).

### Large-scale InDels occur via unequal intra-strand recombination and double-strand break (DSB) repair

To address the underlying mechanisms of large-scale rearrangements, it was important to identify and characterize the InDels breakpoints. Detailed analysis of 9 of the 11 loci (i.e., 3B_1_, 3B_2_, 3B_3_, 3B_4_, 3B_5_, 5B_1_, 5B_2_, 5B_3_, 5B_4_, Table 1) led us to suggest two main mechanisms, namely unequal intra-strand recombination and double strand break repair via non-homologous end-joining (NHEJ).

#### Unequal intra-strand recombination

In 3 of the 9 loci considered (3B_1_, 5B_1_, and 5B_2_; Table 1), high nucleotide identity between the 5’ and 3’ regions of the eliminated sequence was noted. In the 5B_1_ and 5B_2_ loci, the absent sequences from bread wheat genome vs. wild were found to contain sequence duplications, with two direct sequence repeats sharing high nucleotide identity (95% or higher) throughout long sequence segments.

Dot plot comparison of the genomic locus surrounding locus 5B_1_ from wild emmer chromosome 5B and from bread wheat chromosome 5B revealed a 196 kb sequence from wild emmer genome that lacks long segmental similarity to the orthologous locus in bread wheat (Table 1, Supplemental Fig. S2A). This 196 kb segment borders high sequence similarity regions composed of two direct sequence repeats (Supplemental Fig. S3A) and consists of 71.49% TEs. In bread wheat, the 5B_1_ locus is composed of a 7.6 kb segment that shows high nucleotide identity (99%) to both the 5’ flanking sequence (nucleotides 1-1355 and 1798-the end of the locus) and the 3’ flanking sequence (nucleotides 1-1385 and 1798-the end of the locus) of locus 5B_1_ in wild emmer (Fig. 1). The 7.6 kb ^segment from the 5B_1_ locus in bread wheat contains three truncated TEs, *Hawi*, *Clifford* and *Conen*,^ witih ~4 kb in the 3’ region of the segment being annotated as part of a gene coding for lipoxygenase. The InDel in locus 5B_1_ was further validated by PCR analysis using a forward primer based on the 7.6 kb segment in the bread wheat genome, which showed high nucleotide identity to both the 5’ and 3’ regions flanking the wild emmer 5B_1_ locus, and a reverse primer based on the eliminated sequence, which led to wild emmer-specific sequence amplification (Supplemental Fig. S4A). Additional PCR analysis was performed using a forward primer based on the eliminated sequence and a reverse wild emmer-specific primer based on the 3’ flanking region of locus 5B_1_, which showed high nucleotide identity to the 7.6 kb segment in the bread wheat 5B_1_ locus; this also led to wild emmer-specific amplification (Supplemental Fig. S4B). The wild emmer-specific amplification supports bioinformatics-based findings regarding the absence of the 196 kb segment from bread wheat genome, relative to the wild emmer genome. PCR analysis using the same forward primer as used for the reaction portrayed in Supplemental Figure S4A and the reverse primer based on the InDel 3’ flanking region led to amplification of wild emmer and bread wheat sequences (Supplemental Fig. S4C), validating the sequence signature identified at the InDel borders.

**Figure 1.**
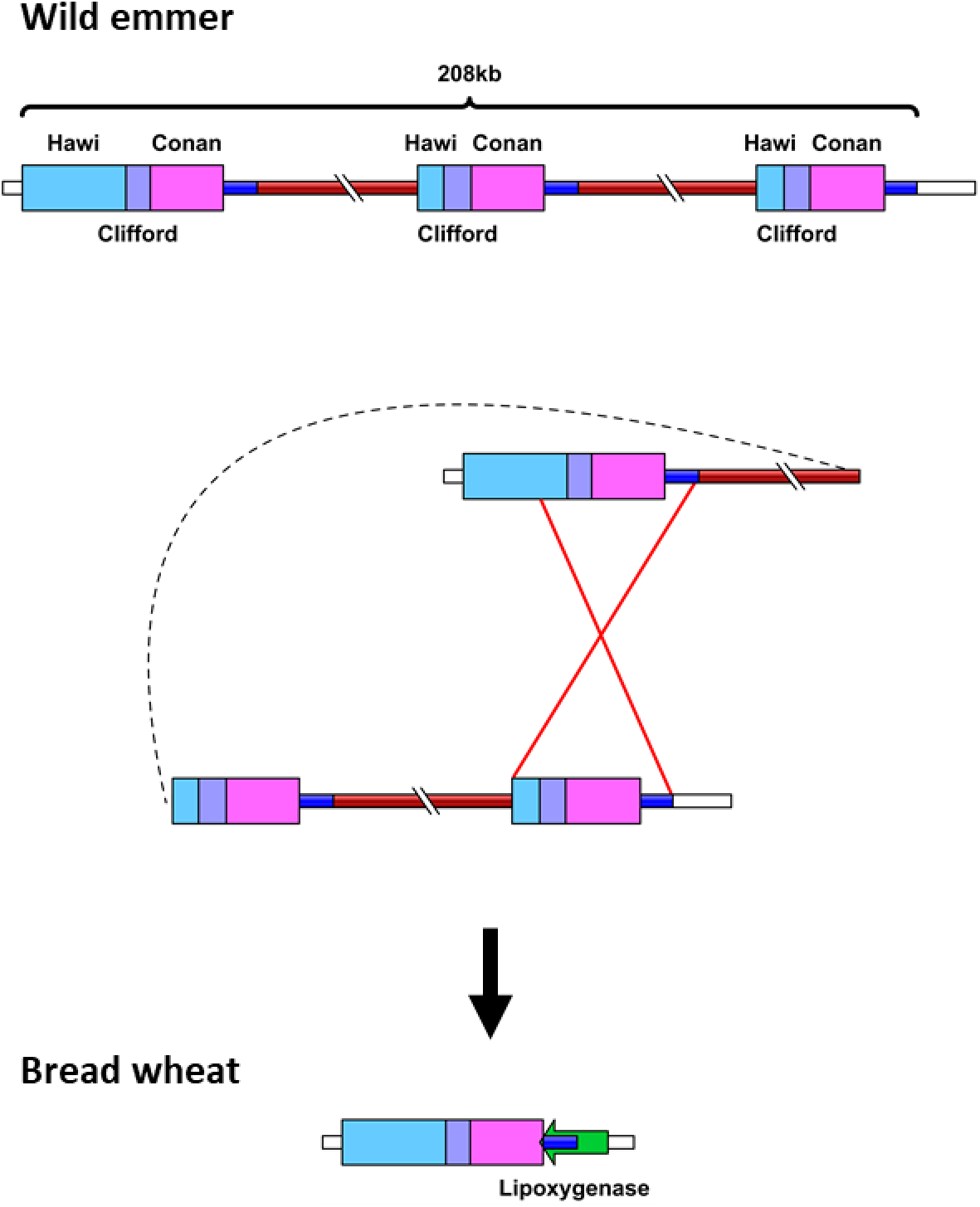
Schematic representation of the locus containing 5B_1_ in the wild emmer (top) and bread wheat (bottom) genomes. Unequal intra-strand recombination involving TEs resulted in a large-scale deletion in bread wheat vs. wild emmer. The *lipoxgenase* gene (green arrow) was annotated in bread wheat, while no genes were identified in the orthologous genomic locus in wild emmer. Sequence length is unscaled. Different colored boxes denote different TE families.

A 252 kb sequence from locus 3B_1_ (Table 1) of wild emmer chromosome 3B was not identified in chromosome 3B of bread wheat. However, the orthologous genomic locus was identified in bread wheat based on sequence alignment between the genomic locus containing 3B_1_ from wild emmer and bread wheat chromosome 3B (Supplemental Fig. S2B). The sequence, which was absent from locus 3B_1_ in bread wheat, was composed of two direct sequence repeats (Supplemental Fig. S3B) and consisted of 61.48% TEs. Locus 3B_1_ in bread wheat consisted of a ~1.8 kb segment, which showed 99% nucleotide identity to the sequence found downstream to 3B_1_ in wild emmer. Additionally, a ~1.5 kb stretch in the 3’ region of locus 3B_1_ in bread wheat showed 92% nucleotide identity to the sequence found upstream of wild emmer locus 3B_1_. The missing sequence data (Ns) ~1.8 kb upstream of the 5’ breakpoint in the wild emmer genome could have interfered with exact determination of the 5’ breakpoint and led to partial alignment of the ~1.8 kb segment to the 5’ flanking end of the InDel. A truncated *XC* element was identified 10 nucleotides downstream of the 5’ end of the 1.8 kb segment in bread wheat and at 10 nucleotides downstream of the locus 3B_1_ 3’ end in wild emmer. An additional truncated *XC* element was annotated 1.1 kb upstream of the 5’ breakpoint of the InDel in locus 3B_1_ in wild emmer.

The TE-containing segments flanking the sequences that were absent in loci 5B_1_ and 3B_1_ in bread wheat vs. wild emmer might have served as a template for unequal intra-strand recombination, resulting in the elimination of the DNA segments between them. Unequal crossing over was recently suggested as being the mechanism involved in the large deletions identified between two allohexaploid wheat cultivars (Thind et al. 2018).

An additional 19 kb sequence consisting of 99.6% TEs was absent in locus 5B_2_ (Table 1) in bread wheat chromosome 5B, relative to wild emmer. The InDel borders were identified using dot plot alignment between the locus containing 5B_2_ in the wild emmer genome and the orthologous locus in bread wheat chromosome 5B. In this manner, the InDel breakpoints were determined as the borders of the high sequence similarity regions (Supplemental Fig. S2C). Both of the InDel breakpoints were located within *Inga* LTRs which share the same orientation, suggesting that this rearrangement might has been the result of sequence elimination due to inter-element recombination, as was previously shown in *Arabidopsis* and rice (Devos et al. 2002; Ma et al. 2004).

#### DSB repair via Non-Homologous End-Joining (NHEJ)

For 6 loci (3B_2_, 3B_3_, 3B_4_, 3B_5_,5B_3_, and 5B_4_) the InDel borders showed only micro-homology (<10 bp), which is not sufficient to serve as a template for homologous recombination (Bennetzen et al. 2005). However, the 6 orthologous loci from which the sequences were eliminated in the bread wheat genome bear sequence signatures characteristic of DSB repair via NHEJ mechanisms. In eukaryotic cells, DSB repair occurs through two main processes, homologous recombination and NHEJ. In plants, DSB repair occurs more frequently via NHEJ than via homologous recombination (Gorbunova and Levy 1997).

NHEJ pathways for DSB repair can be divided as canonical non-homologous end-joining (C-NHEJ) and microhomology-mediated end-joining (MMEJ) processes (Khodaverdian et al. 2017; Ranjha et al. 2018). The C-NHEJ and MMEJ pathways are template independent-mechanisms and thus can generate a wide range of chromosomal rearrangements, including large deletions and template insertions (Gorbunova and Levy 1997; Ceccaldi et al. 2016; Ranjha et al. 2018). DSB repair via C-NHEJ is favored when end resectioning is blocked, instead relying on the repair of blunt-ended breaks or exploiting small microhomologies during the alignment of broken ends (Pannunzio et al. 2014; Ceccaldi et al. 2016). However, when DNA resectioning occurs, other repair pathways, including MMEJ, can compete in repairing the DBS (Ceccaldi et al. 2016). Therefore, DSB repair via MMEJ generates large deletions more often than does DSB repair via C-NHEJ (Ceccaldi et al. 2016; Khodaverdian et al. 2017).

DNA insertions at the DSB repair site, also known as filler DNA, were previously described in plants (Gorbunova and Levy 1997; Salomon and Puchta 1998; Vu et al. 2014; Vu et al. 2017). Filler DNA can be produced when the 3’ ends formed at the break site invade a template, such that synthesis is primed based on a short region of homology. Following one or more rounds of template-dependent synthesis, the newly synthesized DNA can join the second end of the DSB, resulting in template insertion (Gorbunova and Levy 1997; Yu and McVey 2010; Vu et al. 2014; Khodaverdian et al. 2017). The template for filler DNA synthesis seems more often to be found *in cis*, namely on the same molecule, rather than *in trans*, i.e, on another molecule (Wessler et al. 1990; Gorbunova and Levy 1997; Khodaverdian et al. 2017). It was proposed that limited DNA synthesis can lead to the presence of microhomology between the DSB ends, which can then be used for DSB repair via synthesis-dependent microhomology-mediated end-joining (SD-MMEJ) (Gorbunova and Levy 1997; Yu and McVey 2010; Vu et al. 2014; Khodaverdian et al. 2017).

A 64 kb sequence consisting of 79.89% TEs in wild emmer chromosome 5B, locus 5B_3_ (Table 1), was absent in the orthologous genomic locus in the bread wheat genome. However, the orthologous locus from which the 64 kb segment was absent was identified in the bread wheat genome based on flanking alignment. Moreover, the InDel breakpoints were identified by dot plot comparison of the sequences flanking the 5B_3_ locus in the wild emmer and bread wheat genomes (Supplemental Fig. S2D). Locus 5B_3_ was found to border mononucleotide ‘A’ at both the 5’ and 3’ ends in wild emmer, while in bread wheat, the 64 kb segment between the two ‘A’ mononucleotides was absent. Instead, the ‘A’ mononucleotide appeared in a single copy between the conserved sequences flanking locus 5B_3_ and both of the ‘A’ mononucleotides in wild emmer (Fig. 2A). The InDel 5’ breakpoint was identified within the truncated *BARE1* and *WIS* TEs, whereas the 3’ breakpoint was identified within a truncated *Fatima* element. PCR analysis using a forward primer based on the deleted sequence and a reverse primer based on the InDel 3’ flanking region resulted in allotetraploid-specific amplification (Supplemental Fig. S5A). At the same time, PCR amplification using a forward primer based on the InDel 5’ flanking region and the same reverse primer based on the InDel 3’ flanking region led to bread wheat-specific amplification (Supplemental Fig. S5B). These results provide additional support for the InDel identified in the 5B_3_ locus. The fact that allotetraploid-specific amplification was observed using the forward primer directed against a sequence in locus 5B_3_ which was not identified in the orthologous locus in bread wheat could be explained by the absence of the 64 kb segment from locus 5B_3_ in bread wheat. This would prevent amplification in the examined bread wheat accessions. The observed bread wheat - specific amplification using primers based on the InDel flanking sequences suggests that the 64 kb sequence indeed was absent from locus 5B_3_ in bread wheat, resulting in a shorter distance between the surrounding sequences, thus allowing amplification from the bread wheat accessions examined.

**Figure 2.**
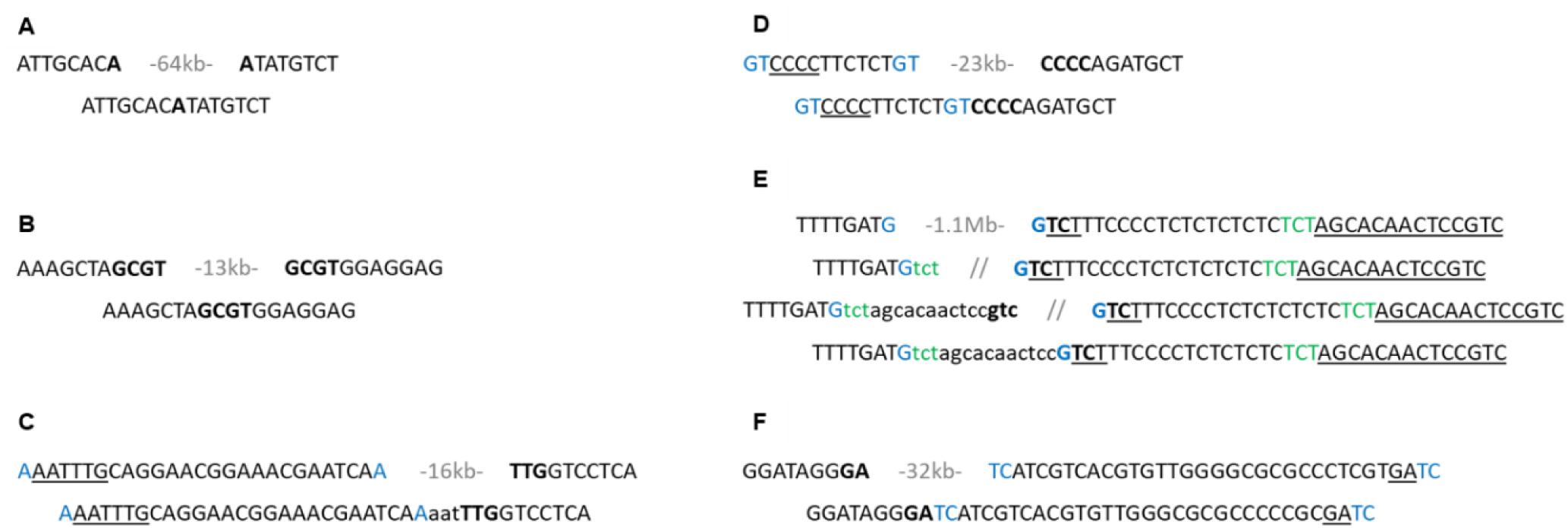
InDels in wild emmer and bread wheat result in sequence signatures characterizing DSB repair via MMEJ (panels A-B) and SD-MMEJ (panels C-F). Sequence signatures from genomic loci 5B_3_ (A), 5B_4_ (B), 3B_2_ (C), 3B_3_ (D), 3B_4_ (E), and 3B_5_ (F). The top row represents the InDel breakpoints in wild emmer, while the bottom row represents the sequence at the orthologous loci in bread wheat. In (E), the second and third rows represent suggested SD-MMEJ intermediates. Only top strands are shown. Bold-short direct or inverted repeats spanning the DSB which might have been utilized for microhomology during DSB repair. Blue and green-short direct repeats near but not necessarily spanning the DSB that might have been used as primer repeats. Templates used in fill-in synthesis are underlined and net sequence insertions are in lowercase. The length of the eliminated sequence is indicated in gray.

In the case of locus 5B_4_ (Table 1), a 13 kb sequence consisting of 81.76% TEs was absent in bread wheat chromosome 5B, as compared to wild emmer. InDel breakpoints were identified by dot plot comparison (Supplemental Fig. S2E), revealing that the 13 kb segment was flanked by the 4-nucleotide motif ‘GCGT’. In bread wheat, a single copy of the ‘GCGT’ motif was identified between the conserved sequences flanking locus 5B_4_ and both of the ‘GCGT’ repeats in wild emmer (Fig. 2B). The 5’ breakpoint was identified within a *Fatima* element.

The InDels identified in loci 5B_3_ and 5B_4_ (Table 1) were flanked by two short tandem repeats (i.e., ‘A’ mononucleotides in locus 5B_3_ (Fig. 2A) and ‘GCGT’ motif in locus 5B_4_ (Fig. 2B)) in wild emmer, while in bread wheat, the sequence between the short tandem repeats was absent and the repeat unit appeared as a single copy. The sequence signature in bread wheat loci 5B_3_ and 5B_4_ was typical for DSB repair via MMEJ, indicating that the InDels in these loci might have resulted from DSB which occurred within the sequences in loci 5B_3_ and 5B_4_ in wild emmer. DSB followed by exonucleases activity and the short tandem repeats that appear in the resulting overhangs could be used for micro-homology in DSB repair via MMEJ. Long deletions with a DSB repair signature similar to that observed in the InDels identified in loci 5B_3_ and 5B_4_ were recently described in two allohexaploid wheat cultivars (Thind et al. 2018).

A 16 kb sequence consisting of 67.63% TEs from locus 3B_2_ (Table 1) in wild emmer chromosome 3B was not identified in bread wheat chromosome 3B. The orthologous genomic locus in the bread wheat genome was identified by alignment of the sequences flanking locus 3B_2_ in the wild emmer genome with bread wheat chromosome 3B. The InDel breakpoints were identified by dot plot comparison (Supplemental Fig. S2F). The 5’ end of the InDel was flanked by the mononucleotide ‘A’, while the 3’ end of the InDel border was flanked by the trinucleotide ‘TTG’, which also appeared 22 bp upstream of the ‘A’ mononucleotide adjacent to the 5’ InDel breakpoint as part of the sequence ‘AAATTTG’ (Fig. 2C). In the bread wheat genome, the 16 kb segment was absent and a trinucleotide template insertion ‘AAT’ was identified between the ‘A’ mononucleotide and the ‘TTG’ trinucleotide. The sequence signature in bread wheat might be the result of DSB repair via SD-MMEJ, whereby following DSB within the 3B_2_ locus and end restriction, the ‘A’ mononucleotide adjacent to the 5’ InDel breakpoint served as a primer repeat, annealed to the first nucleotide in the complementary strand of the sequence ‘AAATTTG’ found upstream of the InDel 5’ breakpoint, thus enabling the synthesis of the 6-nucleotide ‘AATTTG’. This synthesis led to trinucleotide (’TTG’) micro-homology between the right and left sides of the break, which was used for annealing, and resulted in an InDel junction including a trinucleotide insertion (’AAT’) and deletion of the 16 kb segment from the 3B_2_ locus (Fig. 2C). The InDel 5’ breakpoint was identified within the truncated TE *Mandrake* and the 3’ breakpoint was identified within an intact *Fatima* element.

An additional 23 kb segment from locus 3B_3_ (Table 1) in wild emmer consisting of 99.55% TEs was not identified in bread wheat chromosome 3B. However, the orthologous genomic locus from bread wheat was identified by flanking alignment, while the InDel breakpoints were determined by dot plot comparison of the locus flanking locus 3B_3_ in the wild emmer and bread wheat genomes (Supplemental Fig. S2G). Locus 3B_3_ in the bread wheat genome carries the signature of DSB repair via SD-MMEJ (Fig. 2D). The 5’ breakpoint of the InDel in locus 3B_3_ borders with the dinucleotide ‘GT’. Additional ‘GT’ dinucleotide motif appeared as a tandem repeat 12 bp upstream of the ‘GT’ dinucleotide adjacent to the 5’ breakpoint, followed directly by the 4-nucleotide ‘CCCC’ motif. The 3’ breakpoint of the InDel identified of locus 3B_3_ was also bordered by a ‘CCCC’ motif. The dinucleotide ‘GT’ might thus have been used as a primer repeat, thereby enabling the synthesis of the ‘CCCC’ motif. DSB via SD-MMEJ resulted in the generation of an apparently blunt repair junction and elimination of the 23 kb segment. Alternatively, the blunt repair junction observed could be the result of DSB repair via C-NHEJ. However, the long deletion suggests that following the DSB, DNA resectioning based on exonuclease activity occurred. As such, DSB repair via MMEJ is more likely to have occurred (Ceccaldi et al. 2016). Finally, the InDel 5’ breakpoint was identified within a truncated *Xalax* TE and the 3’ breakpoint was identified within an intact *Fatima* element.

A 1.1 Mb sequence in the wild emmer 3B_4_ locus (Table 1) consisting of 77.64% TEs was found to border mononucleotide ‘G’ and was not identified within bread wheat chromosome 3B (Fig. 2E). However, the orthologous locus was identified in the bread wheat genome based on flanking alignment. The InDel breakpoint was identified by dot plot comparison of the genomic region containing the 3B_4_ locus in wild emmer chromosome 3B and in bread wheat chromosome 3B (Supplemental Fig. S2H). The InDel in locus 3B_4_ resulted in a 14 bp templet insertion into the bread wheat genome (’TCTAGCACAACTCC’), bounded by ‘G’ mononucleotides, which formed a direct repeat with a sequence found 20 bp downstream of the ‘G’ mononucleotide adjacent to the 3’ breakpoint in wild emmer (Fig. 2E). This InDel junction could have arisen as a result of DSB repair that included two rounds of *trans* microhomology annealing and synthesis. In this scenario, during the DSB repair which occurred between the wild emmer and bread wheat genomes, the ‘G’ mononucleotide found at the 5’ end of the DSB served as a primer repeat and annealed to the nucleotide complimentary to the ‘G’ mononucleotide found in the 3’ end of the DSB, thus enabling synthesis of the trinucleotide ‘TCT’. The newly synthesized ‘TCT’ motif at the 5’ end of the DSB was then annealed to the complimentary sequence of the ‘TCT’ trinucleotide found 20 bp downstream of the ‘G’ mononucleotide adjacent to the 3’ InDel breakpoint in wild emmer, thus resulting in the synthesis of the sequence ‘AGCACAACTCCGTC’. Following two rounds of nucleotide synthesis, trinucleotide (’GTC’) microhomology between the right and left sidez of the break used for annealing resulted in an InDel junction including a 14 bp templated insertion and the elimination of 1.1 Mb sequence. Additionally, a variation in the copy numbers of the dinucleotide ‘TC’ repeat found 9 bp downstream of the 3’ breakpoint was identified in wild emmer (7 tandem repeats of the dinucleotide) and bread wheat (6 tandem repeats of the dinucleotide) was observed. The InDel 5’ breakpoint was identified within a truncated *Egug* TE. The deleted sequence included a gene of unknown function and a gene coding for an uncharacterized protein. Additional support for the sequence elimination from locus 3B_4_ was obtained upon PCR analysis using primers based on the InDel flanking sequences and on the deleted sequence (Supplemental Fig. S6A-B). PCR analysis using a forward primer based on the 5’ flanking sequence of locus 3B_4_ and a reverse primer based on the deleted sequence yielded an emmer-specific amplification (Supplemental Fig. S6A). At the same time, PCR using the same forward primer and a reverse primer designed from the 3’ flanking region of locus 3B_4_ resulted in amplification in the bread wheat accessions examined but no amplification in wild emmer.

An additional 32 kb sequence consisting of 98.22% TEs from the 3B_5_ locus (Table 1) in wild emmer chromosome 3B was absent in the orthologous locus in the bread wheat genome. The conserved sequences flanking the 3B_5_ locus in bread wheat, identified by dot plot alignment (Supplemental Fig. S2I), were found to connected by an apparent blunt end junction. In wild emmer, the 3B_5_ locus bordered with the dinucleotide ‘GA’ at the 5’ end and with the dinucleotide ‘TC’ at the 3’ end. A 4-nucleotide ‘GATC’ motif was found 29 bp downstream of the dinucleotide ‘TC’ adjacent to the 3’ breakpoint of the InDel (Fig. 2F). The sequence signature in the bread wheat 3B_5_ locus corresponded to a site of DSB repair via SD-MMEJ, with the ‘GA’ motif on the complimentary strand to the dinucleotide ‘TC’ found at the 3’ breakpoint serving as a primer repeat used for annealing to the 4-nucleotide ‘GATC’ motif found 29 bp downstream ofthe ‘TC’ dinucleotide adjacent to the 3’ breakpoint, thus enabling synthesis of ‘TC’ dinucleotide on the complimentary strand from the 3’ end of the DSB. In this scenario, dinucleotide synthesis led to the appearance of dinucleotide (’TC’) microhomology between the DSB ends, which were then annealed to yield the apparent blunt end junction seen in the bread wheat genome. The apparent blunt ends junction may also be the result of DSB repair via C-NHEJ. However, repair via C-NHEJ is less likely, considering the length of the eliminated sequence. Four SNPs were detected in the sequence found 17-35 bp downstream of the ‘TC’ dinucleotide adjacent to the 3’ breakpoint in wild emmer. The InDel breakpoints were found within intact (5’ breakpoint) and truncated (3’ breakpoint) *Fatima* elements.

### Introgression of DNA fragments of unidentified origin into the wheat genome

The InDel in chromosome 5B locus 5B_5_ (Table 1) was revealed based on sequence alignment of the flanking sequences of a wild emmer-specific *Fatima* insertion into bread wheat chromosome 5B. Following the identification of the orthologous locus in bread wheat chromosome 5B, the InDel breakpoints were determined as the borders of the gaps observed in both axes by dot plot comparison of the orthologous loci from the wild emmer and bread wheat genomes (Supplemental Fig. S2J). The InDel identified in locus 5B_5_ involved the replacement of a 41 kb segment consisting of 98.18% TEs found in the wild emmer genome with a 11 kb segment consisting of 61.48% TEs located in the orthologous genomic locus in the bread wheat genome (Fig. 3). The InDel locus 5B_5_ 5’ breakpoint was found within a truncated *Karin* TE, while the 3’ breakpoint was found within a truncated *Deimos* TE. PCR validation was carried out using primers based on the flanking sequences of the InDel coupled with primers designed against the 41 kb wild emmer-specific segment, resulting in wild emmer-specific amplification (Supplemental Fig. S7A-B). The third PCR amplification used a forward primer based on the 11 kb bread wheat-specific sequence and the same reverse primer based on sequence located downstream to the InDel, as used in the previously described reaction. This third PCR resulted in amplification of both of the examined bread wheat accessions, yet no amplification was observed for wild emmer (Supplemental Fig. S7C).

**Figure 3.**
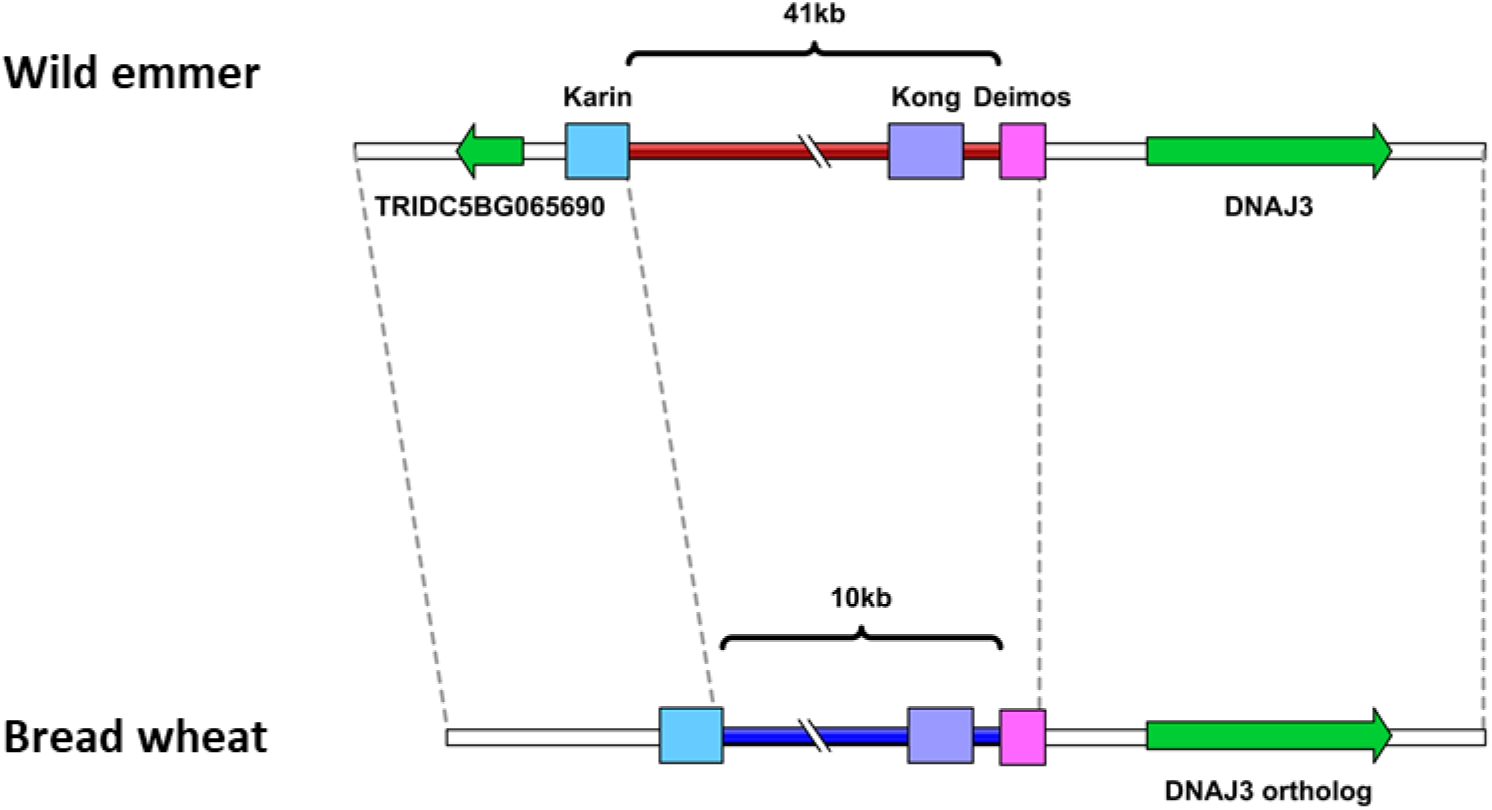
Schematic representation of locus 5B_5_ in wild emmer (top) and bread wheat (bottom). Introgression of a new sequence into locus 5B_5_ in the wheat genome. Sequence length is unscaled. Colored boxes denote different TE families. Genes are represented by green arrows.

The 11 kb sequence in bread wheat locus 5B_5_ was not identified within the wild emmer or *Ae. tauschii* (the donor of the D sub-genome) genomes based on sequence alignment. A possible explanation for this phenomena is introgression of a new sequence into the wheat genome, namely the transferring of DNA segments from one species into another via recurrent backcrossing (Mirzaghaderi and Mason 2017). Indeed, introgression of chromosomal segments from alien genomes is known to be facilitated by allopolyploidy in the wheat group (Feldman and Levy 2012).

### Variations in copy numbers of a long tandem repeat in wild emmer vs. bread wheat

The analysis of locus 5B_6_ (Table 1) in chromosome 5B revealed variations in the copy numbers of a ~460 kb segment, which appeared as two tandem repeats in the wild emmer genome (totaling 924 kb in length and comprising 79.57% TEs) and in a single copy (422 kb in length and comprising 78.04% TEs) in the bread wheat genome (Fig. 4). This copy number variation was identified by dot plot comparison of the orthologous locus surrounding locus 5B_6_ in wild emmer and bread wheat (Supplemental Fig. S2K). The 422 kb segment in locus 5B_6_ in bread wheat showed high sequence similarity (95% or higher with a word size of 100 through long sequence segments) to two repeat units observed in the orthologous locus in wild emmer. The borders of the single repeat unit in bread wheat were determined based on discontinuity points in the sequence coverage (Supplemental Fig. S2K). The borders of the tandem repeats in wild emmer were determined by dot plot comparison of the locus ^surrounding locus 5B_6_ in wild emmer against its self, as the borders of the regions showing high^ sequence identity through long sequence segments (95% or higher with a word size of 100) outside of the diagonal line represent the continuous match of the sequence to its self (Supplemental Fig. S3C). In wild emmer, a gene coding for an F-box domain-containing protein was annotated 176 bp downstream of the 5’ end of the first repeat, while a gene of unknown function was annotated to the 3’ end of the second repeat. Additionally, the first repeat in wild emmer contained a gene coding for the coatomer beta subunit. In bread wheat, the 3’ end of the single repeat was identified within a protein coding gene and three additional high confidence protein coding genes were identified within the sequence that underwent copy number variation. The genomic locus in which locus 5B_6_ was found underwent inversion between wild emmer and bread wheat. The borders of the inversion were identified and the inversion length was determined to be ~6.5 Mb (Supplemental Fig. S2L).

**Figure 4.**
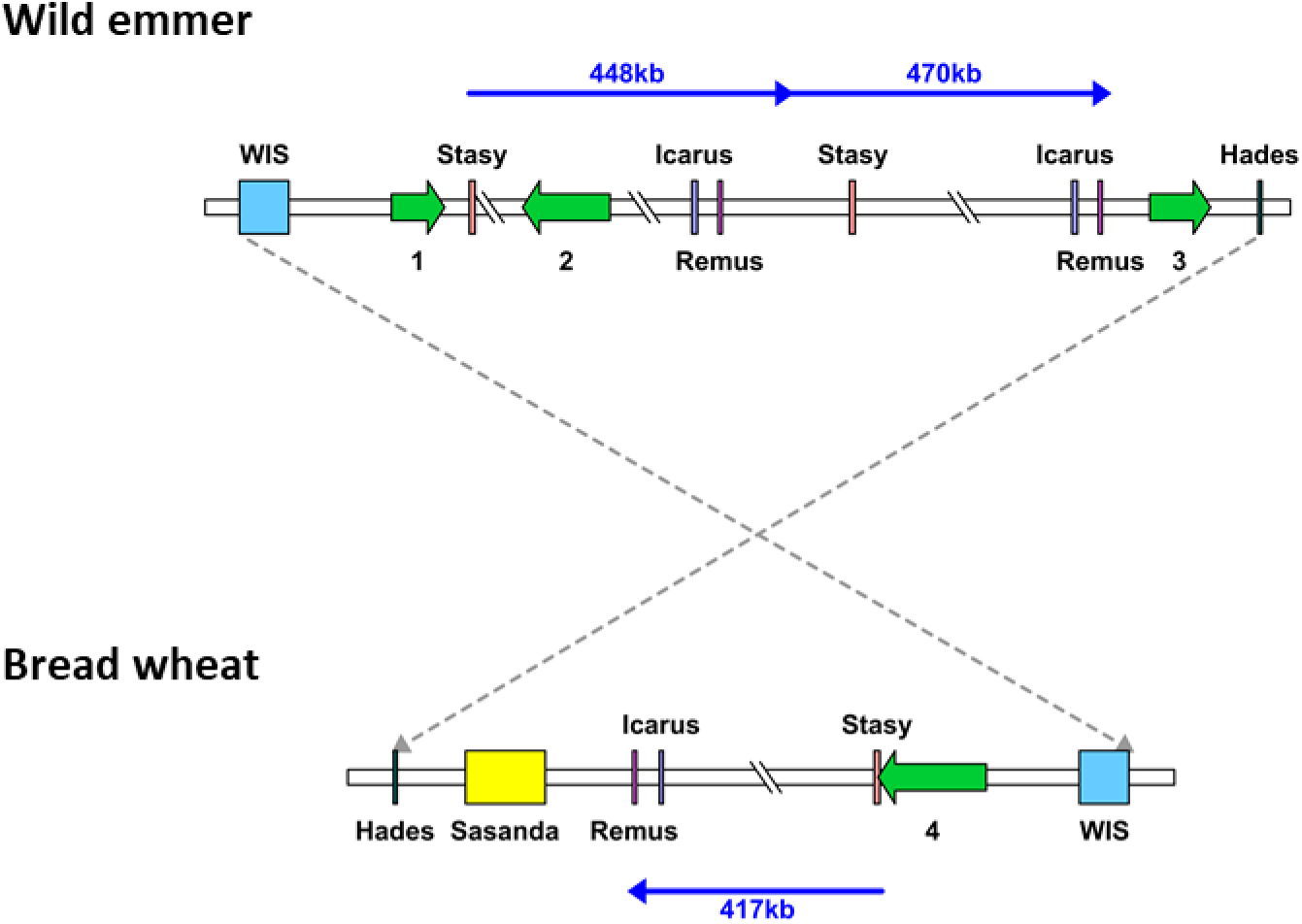
Schematic representation of locus 5B_6_ in wild emmer (top) and bread wheat (bottom). Segmental duplication in wild emmer locus 5B_6_. Sequence length is unscaled. Locus 5B_6_ is part of a ~6.5 Mbp segment that underwent inversion between wild emmer and bread wheat. TEs are represented as colored boxes. Genes are denoted by green arrows: 1. TRIDC5BG011160.1, F-box domain-containing protein; 2. TRIDC5BG011170.1, Coatomer, beta subunit; 3. TRIDC5BG011180, unknown function; 4. TRIAE_CS42_5BS_TGAC_424303_AA1388580, Protein coding.

The copy number variation observed can be explained by elimination of one of the repeats from the bread wheat genome through unequal intra-strand recombination. Alternatively, the copy number variation seen could be the result of a duplication that occurred within the wild emmer genome later during evolution. To identify the source of the copy number variation and to estimate when this copy number variation transpired, it was important to estimate the numbers of copies of the tandem repeat within different accessions of wheat allopolyploids (3 wild emmer accessions, 3 durum accessions and 4 bread wheat accessions) and within the available species that are closely related to the diploid B sub-genome donor (3 *Aegilops speltoides* accessions and 3 *Aegilops searsii* accessions). The presence of a single repeat was verified by PCR using a forward primer designed against the 5’ flanking region (in the wild emmer genome) of the sequence that underwent copy number variation and a reverse primer designed against the 5’ region of the repeat unit (in the wild emmer genome). Amplification was observed in all the tested accessions of *Ae. speltoides*, wild emmer, durum and *T. aestivum*, suggesting that the examined segment exists in at least one copy in each of these species (Supplemental Fig. S8A). No amplification was observed for the tested *Ae. searsii* accessions (Supplemental Fig. S8A). To examine whether the ~460 kb segment appears as a tandem repeat in the different accessions, PCR was performed using a forward primer based on the sequence located at the 3’ end (in the wild emmer genome) of the segment that underwent copy number variation and the reverse primer that was used in the previously mentioned reaction. The observed *Zavitan*-specific amplification (Supplemental Fig. S8B) suggests that the ~460 kb segment is found as a tandem repeat only in this accession, out of the 16 accessions examined. The PCR results, together with the high sequence identity between the repeats in wild emmer (Supplemental Fig. S3C), support a scenario iwhereby copy number variation is the result of a recent duplication in wild emmer. The boundaries of new segmental duplications in humans were found to be enriched in *Alu*-SINE elements, indicating a possible role for SINE elements in the duplication event (Bailey et al. 2003; Jurka et al. 2004). The presence of a truncated *Stasy* element (SINE) 2.5 kb downstream of the first repeat start in wild emmer and of a highly similar (99%) truncated *Stasy* element in the 5’ region of the second repeat could indicate a possible role for this element in the copy number variation reported here.

### The dynamic nature of the wheat genome

Allopolyploidization involves Inter-generic hybridization and chromosome doubling, two "genetics shocks" which induce rapid genomic changes in the new allopolyploid (Feldman and Levy 2005). In addition to rapid, revolutionary genomic changes, allopolyploid species experience relatively faster evolutionary changes due to mutation buffering in the polyploid genome (Van de Peer et al. 2017). Allopolyploid genome plasticity, together with heterosis, may play an important role in the success of some allopolyploid lineages, relative to their diploid progenitors (Feldman and Levy 2012; Van de Peer et al. 2017). Large-scale genomic rearrangements between wheat allopolyploids (Devos et al. 1995; Badaeva et al. 2007; Jorgensen et al. 2017; Huang et al. 2018; Thind et al. 2018) and between wheat allopolyploids and their progenitor species (Dvorak et al. 2018; Huo et al. 2018) were previously identified. However, the mechanisms involved in such rearrangements remain largely unknown.

Domestication, together with allopolyploidization, is a key event shaping the wheat genome through selection (Avni et al. 2017; Akpinar et al. 2018). To better assess when the structural variations identified in this study occurred, site-specific PCR analyses were performed for 4 tested accessions, namely a sequenced accession of wild emmer (*Zavitan*), an accession of durum (*Svevo*) and two accessions of bread wheat (accessions CS46 and TAA01). Primers used were based on five of the sequence variations identified in this paper, as described previously (Supplemental Figs. S4-S8). For the InDel of locus 5B_3_, similar amplification patterns from the tested wild emmer and durum accessions (Supplemental Fig. S5) suggested this InDel occurred following allohexaploidization or during hexaploid wheat evolution. However, for the InDels of loci 5B_1_, 3B_4_ and 5B_5_, the similar amplification patterns seen for durum and bread wheat (Supplemental Figs. S4, S6 and S7) indicated that these InDels occurred during the evolution of tetraploid wheat, possibly during wheat domestication. The availability of a high-quality durum genome assembly will allow for better characterization of the evolutionary time frame and the events leading to genomic rearrangements in wheat.

TEs are a great source for mutations not only due to their repetitive nature which makes them a target for homologous recombination but also as a direct result of their actions. Besides simple insertion or element excision, TE activity might trigger DSBs at insertion and excision sites (Gray 2000; Hedges and Deininger 2007; Wicker et al. 2010). Additionally, alternative transposition events can also result in TE-associated chromosomal rearrangements (Gray 2000). The high TE content of the wheat genome, together with the high genome plasticity that characterizies polyploid genomes, could contribute dramatically to the diversity observed among wheat allopolyploids. In the present study, previous knowledge of how elimination of *Fatima*-containing sequences following allopolyploidization may have contributed to the relative high efficiency of our analysis. Following manual data validation, only 4% of the polymorphic insertion sites were removed from the analysis as they were most likely the result of assembly artefacts (missing sequencing data –N_s_--in one or both of the identified breakpoints).

In summary, we suggest that sequence deletions mediated through DSB repair and unequal intra-strand recombination, together with the introgression of new DNA sequences, contribute to the large genetic and morphological diversity seen in wheat allopolyploids and to their ecological success, relative to their diploid ancestors. Such large-scale genomic rearrangements are most likely facilitated by allopolyploidization. The presence of TEs in InDels borders suggests a possible role for TEs in the large-scale genomic rearrangements seen in wheats allopolyploids, either acting via homologous recombination or other mechanisms. Accordingly, this study aimed to uncover the underlying mechanisms of DNA elimination in wheat, a phenomenon that remained unsolved for many years. Better assembly of the wheat genome drafts will allow for assessing the extent of large-scale DNA rearrangements and evaluating their impact on genome size.

## Materials and methods

### Plant material and DNA isolation

In this study, we used 17 accessions of *Triticum* and *Aegilops* species (Supplemental Table S1): 3 wild emmer (*T. turgidum* ssp*. dicoccoides*) accessions, including the sequenced accession *Zavitan*; 3 durum (*T. turgidum* ssp*. durum*) accessions, including *Svevo*; 4 bread wheat accessions, including two *Chinese Spring* accessions (CS46 and TAA01); six B genome diploid accessions (*Ae. speltoides*-3 accessions, *Ae. searsii*-3 accessions) and a single *Ae. tauschii* accession. DNA was extracted from young leaves ~4 weeks post-germination using the DNeasy plant kit (Qiagen, Hilden, Germany).

### Wheat genomic data

The genome drafts of three *Triticum* and *Aegilops* species were used in this study: (1) WEW (wild emmer wheat) assembly, a full genome draft of emmer wheat that was sequenced using paired-end and mate-pair shotgun sequencing and assembled using DeNovoMAGIC. The WEW assembly (http://wewseq.wix.com/consortium) contains sorted chromosomes and covers ~95% of the emmer wheat genome (Avni et al. 2017). (2) The bread wheat *T. aestivum* Chinese Spring assembly (downloaded in June, 2017 from http://plants.ensembl.org/Triticum_aestivum/Info/Index) was generated by the International Wheat Genome Sequencing Consortium (IWGSC). This assembly covers 14.5 Gbp of the genome with an N50 of 22.8 kbp. Pseudomolecule sequences were assembled by integrating a draft *de novo* whole-genome assembly (WGA), based on Illumina short-read sequences using NRGene deNovoMagic2, with additional layers of genetic, physical, and sequence data (Appels et al. 2018). (3) The Aet v4.0 assembly, a reference quality genome sequence for *Ae. tauschii* ssp. *strangulate* (data available from the National Center for Biotechnology Information (NCBI)), was generated using an array of advanced technologies including ordered-clone genome sequencing, whole-genome shotgun sequencing and BioNano optical genome mapping and covers 4.2 Gbp of the genome (Luo et al. 2017).

### Retrieving *Fatima* insertions froHT m wild emmer and bread wheat draft genomes

A specific variant of intact *Fatima* element and flanking sequences (500 bp from each side) were retrieved from wild emmer and bread wheat draft genomes using MITE analysis kit (MAK) software (http://labs.csb.utoronto.ca/yang/MAK/) (Yang and Hall 2003; Janicki et al. 2011). The publicly available consensus sequence of the *Fatima* element RLG_Null_Fatima_consensus-1 (9997 bp in length) was downloaded from TREP database (http://wheat.pw.usda.gov/ggpages/Repeats/) and used as input (query sequence) in the MAK software. BLASTN was performed against the draft genomes. For retrieval of the *Fatima* sequences, the MAK “member” function was used with an e-value of e^−3^ and an end mismatch tolerance of 20 nucleotides. In addition, flanking sequences (500 bp from each end) were retrieved together with each of the *Fatima* insertions to characterize insertion sites.

### Identification of species-specific *Fatima* insertions

To identify potentially species-specific *Fatima* insertions, the flanking sequences of the retrieved *Fatima* elements from the wild emmer 3B and 5B chromosomes were aligned to the flanking sequences of thosee elements retrieved from the orthologous chromosomes in bread wheat. Alignments were performed with BLAST+ stand-alone version 2.2.24, using an e-value less than e^−100^. *Fatima* elements in wild emmer for which no flanking similarity was identified in the orthologous bread wheat chromosome were considered as candidate wild emmer-specific insertions and were further examined. Additionally, a case where two *Fatima* insertions from the wild emmer genome showed high flanking similarity to a single *Fatima* insertion from the bread wheat genome was examined.

### Identification and characterization of *Fatima*-containing sequences that undergo InDel and of InDel breakpoints

The flanking sequences of the candidate wild emmer-specific insertions were compared to bread wheat chromosome 3B or 5B, depending on the insertion location in wild emmer, using BLAST to identify the orthologous genome locus. In cases where the orthologous genome locus has yet to be identified, a chromosome walking approach was employed, such that longer flanking sequences of the *Fatima* insertion in wild emmer were aligned to the orthologous chromosome from the bread wheat genome using BLAST. Following identification of the orthologous genome locus, dot plot alignments, corresponding to graphical representations of sequence aliments, were performed on orthologous loci to identify sequence variations, using UGENE version 1.23.0 (Okonechnikov et al. 2012) with a minimum repeat length of 100 bp and 95% repeat identity. For each InDel observed, the sequence alignments were analyzed and the breakpoints, namely regions where sequence similarity broke down, were identified. To determine InDel lengths, the distance between two breakpoints was calculated, based on a minimum repeat length of 100 bp and 95% repeat identity.

To further characterize InDels, breakpoints and deleted and inserted sequences were annotated to genes and TEs. Gene annotation was performed using The Grain-Genes Genome Browsers (https://wheat.pw.usda.gov/GG3/genome_browser) for wild emmer and bread wheat and the *EnsemblPlants* (http://plants.ensembl.org/Triticum_aestivum/Info/Index) genome browser for bread wheat. TE annotation was performed using Repeat-Masker (http://www.repeatmasker.org/) with a cutoff of 250 and TE databases of wheat transposable elements taken from TREP (http://wheat.pw.usda.gov/ggpages/Repeats/).

### PCR analysis

PCR validation was performed using primers designed with PRIMER3 version 4.1.0 based on identified sequence variations (see Supplemental Table S2 for primer sequences), such as f eliminated or newly introduced sequences and sequences flanking eliminated segments. To generate PCR products up to 800 bp, each reaction contained: 10µl PCRBIO HS Taq Mix Red (PCRBiosystems), 7 µl ultrapure water (Biological Industries), 1 µl of each site-specific primer (10µM) and 1 µl of template genomic DNA (approximately 50 ng/µl). The PCR conditions were 95°C for 2 min, 35 cycles of 95°C for 10 sec, the calculated annealing temperature for 15 sec and 72°C for 15 sec. For PCR products longer than 800 bp, each reaction contained 12 µl ultrapure water (Biological Industries), 4 µl of 5X PrimeSTAR GXL Buffer (TaKaRa), 1.6 µl of 2.5 mM dNTPs, 0.5 µl of each site-specific primer (10 µM) and 0.4 µl of PrimeSTAR GXL DNA Polymerase (1.25 U, TaKaRa). The PCR conditions used were 94°C for 5 min, 30 cycles of 98°C for 10sec, the calculated annealing temperature for 15 sec and 68°C for 1 min. PCR products were visualized in 0.8-1% agarose gels.

## Acknowledgements

We want to thank Dr. Guojun Yang, University of Toronto, for providing the stand-alone version of MAK. We also thank Prof. Avi A. Levy, Weizmann Institute of Science, and Prof. Assaf Distelfeld, Tel-Aviv University, for their critical reading of the manuscript. This work was supported by a grant from the Israel Science Foundation (322/15) to K. K.

## Author contributions

**Inbar Bariah**: Performed the research, analyzed the data, wrote the paper.

**Danielle Keidar-Friedman**: Performed the research, analyzed the data, wrote the paper.

**Khalil Kashkush**: Corresponding author, designed the research, analyzed the data, wrote the paper.

